# CD271 sorting for improved liver cell isolation: Semiautomated and simultaneous preparation of parenchymal and non-parenchymal cells from mouse and human livers

**DOI:** 10.1101/2025.01.21.633876

**Authors:** Anne Dropmann, Bedair Dewidar, Kerry Gould, Raquel L. Baccetto, Christoph Meyer, Tiziana Caccamo, Pia Erdoesi, Şamil Ayvaz, Andrea Scheffschick, Georg Damm, Daniel Seehofer, Laura Kim Feiner, Bianca Kruse, Claudia Rubie, Matthias Glanemann, Vanessa Orth, Emrullah Birgin, Nuh Rahbari, Matthias P. Ebert, Steven Dooley, Seddik Hammad

## Abstract

**Background:** A detailed understanding of the dynamic fate changes of hepatocytes, hepatic stellate cells (HSC), Kupffer cells (KC), and liver sinusoidal endothelial cells (LSEC) is critical for studying liver (patho)physiology during disease progression. Current isolation methods often focus on single cell types, limiting utility in comprehensive research.

**Aim:** To develop a novel, semi-automated protocol for the simultaneous isolation of hepatocytes and non-parenchymal cells (NPCs), including HSC, KC, and LSEC, from mouse and human, with high yield, purity, and viability from healthy and diseased livers.

**Method:** The protocol employs a two-step EGTA and collagenase II perfusion for tissue digestion. Hepatocytes were isolated by low-speed centrifugation and a Percoll gradient. Subsequently, magnetic-activated cell separation, using CD271 as a selective surface marker for HSC, CD11b for KC and CD146 for LSEC) was performed. Validation was achieved with immunofluorescence staining, flow cytometry, RT-PCR, and UV fluorescence, whereby yield, purity, and viability were assessed.

**Results:** With our method, yield of hepatocytes, HSC, KC, and LSEC, is 33.4±5.5×10⁶, 5.2±6.3×10⁴, 12.4±4.8×10⁵ and 18.2±8.9×10⁵ cells per healthy mouse liver, respectively, with cell viabilities exceeding 89%, and purity surpassing 90%. CD271 was validated as an effective marker for purifying HSC in healthy and diseased human (n=4-6) and mouse livers. Compared to microfluidic and organ-on-a-chip approaches, with our protocol, we achieve higher yield and purity values while enabling the simultaneous isolation of multiple cell types from a single sample.

**Conclusion:** Our semi-automated protocol offers a scalable, reliable, and versatile solution for isolating main liver cell types with high yield, purity, and viability from both healthy and diseased tissues, advancing liver research and facilitating downstream investigations.

**Impact and implications:**

1. **Broad applicability**: The CD271-based method efficiently isolates key liver cell types (Hepatocyte, HSC, KC, LSEC) simultaneously.
2. **Versatility in disease models**: Effective for studying healthy, fibrotic, and damaged liver tissues.
3. **Robust across variability**: Works reliably across different mouse strains, age groups, and conditions.
4. **Human research potential**: Scalable for high-purity isolation from human liver tissue, enabling translational studies.
5. **High-quality results**: Ensures >85% viability and >90% purity, supporting reproducible liver research applications.

## Introduction

The liver is a multi-functional and multi-cellular organ with diverse functions, playing a key role in maintaining homeostasis and adapting to pathological challenges. Hepatocytes play a central role by regulating nutrient metabolism, biotransformation, and synthesizing key systemic proteins such as albumin and coagulation factors, alongside bile acid production and transport. These aforementioned Hepatocyte functions are supported by non-parenchymal cells (NPC), including hepatic stellate cells (HSC), Kupffer cells (KC), and liver sinusoidal endothelial cells (LSEC), which together coordinate liver responses to challenges like toxic injury (1, 2). In humans, hepatocyte comprise approximately 80% of liver cells, while HSC account for 1.4%, KC for 2%, and LSEC for 2.5%, with the remaining cells including cholangiocytes, vascular endothelial cells (3), and pit cells (4), the liver-specific natural killer cells. These cell types in concert maintain liver homeostasis, display a high plasticity and contribute substantially to the different liver responses of the liver to injury.

Liver cells are invaluable for investigating molecular mechanisms underlying diseases such as hepatitis, fibrosis, and cancer, as well as for drug development and toxicity studies. Primary liver cells provide distinct advantages over immortalized cell lines, as they more accurately mimic *in vivo* conditions as suggested by Gomez-Lechon and co-workers (5). Recent advancements, including 3D culture systems (6), organoids (7), precision slices (8), or flow bioreactors (9), have improved experimental models of liver physiology and pathology. However, the availability of primary cells remains a significant bottleneck, with isolation methods requiring substantial time, technical expertise, and access to living animals or well-organized organ resection procedures. Critical parameters such as yield, viability, and purity directly influence the quality of downstream applications.

Traditional liver cell isolation methods typically focus on a single cell type. For instance, hepatocytes are often isolated through low-speed centrifugation and further purified with density gradient, while a particular cell type i.e. LSEC is separated using CD146 surface marker. Although these methods have been refined for single cell type and specific applications (10, 11), they are generally optimized for healthy tissues, where cell phenotypes are well defined. Diseased tissues, however, present unique challenges, as pathological conditions can alter cell properties such as density, morphology, and marker expression, which complicate the isolation protocols.

Simultaneous isolation of multiple liver cell types from a single specimen has proven challenging. For example, Werner and co-workers (10) employed collagenase perfusion, low-speed centrifugation, and magnetic-activated cell sorting (MACS) to purify hepatocytes and NPCs from human liver tissue. Their method yielded high-quality cells but required large tissue samples (25–100 grams), which are not always feasible. Stradiot and co-workers (12) and Zhou et al. (13) developed a fluorescence-activated cell sorting (FACS)-based method to isolate NPCs using scavenging activity markers for LSEC i.e. CD32, (12) but this technique may present challenges that reduce its efficacy or reliability in human cell isolation. These methods emphasize the need for robust, semi-automated or fully-automated, and flexible cell isolation protocols that are independent of tissue volume, disease states, age, and species.

Conventional protocols for HSC isolation rely on density gradient centrifugation following collagenase digestion. However, activated HSC, which represent a major cell fraction in disease states, exhibit altered densities, complicating their purification. An emerging solution involves using specific surface markers for HSC, enabling refinement of protocols for all HSC. Among these surface markers, nerve growth factor receptor (namely CD271, NGFR or p75NTR) has been shown as an HSC-specific marker. Expressed on murine and human HSC in both healthy and diseased livers, CD271 offers a reliable target for isolating HSC, as demonstrated by (14) and (15). Recent studies on CD271 in the liver, particularly in the context of HSC, highlight its role in identifying functional populations (16). In this study, single-cell RNA sequencing has been used to analyze the gene expression profiles of HSC, both in healthy and diseased states (16).

Building on these insights, we have developed a semi-automated pipeline for the simultaneous isolation of hepatocytes, HSC, KC, and LSEC from mouse and human liver specimens. This protocol utilizes CD271 for HSC separation and combines MACS technology with optimized digestion. Our method achieves high yields, purity, and viability across all cell types, irrespective of species, age, strain, or disease stage. For example, the protocol reliably isolates quiescent and activated HSC from both healthy and diseased mouse livers. Additionally, it is adaptable and extendible, allowing the incorporation of novel markers to target other liver cell subtypes.

## Material & Methods

### Mouse models and treatment

Balb/C and C57BL/6 mice (1.5–2, 6–8, and more than 18 months old) were purchased from Janvier Labs (France). Mice were kept under specific pathogen-free conditions in a fixed 12-hour light/dark cycle with free access to normal chow and water *ad libitum*. Additionally, to induce acute liver damage, 6–8-month-old Balb/C mice were exposed to a single i.p. injection of CCl₄ (320 mg/kg, Sigma-Aldrich, Cat. no. 319961). Four-month-old Balb/C mice with a defective Mdr2 bile transporter (Mdr2*KO* mice) were used as a model for chronic biliary-derived liver fibrosis. To investigate toxic-induced fibrosis, adult male C57Bl/6 mice received 1.6 g/kg body weight of CCl₄ intraperitoneally twice per week for 6 weeks (17). Genotyping of the Mdr2*KO* mice was done as previously described (18). All animal experiments followed international guidelines and received prior approval from the local regulatory authorities.

### Simultaneous isolation of hepatocytes and non-parenchymal liver cells from mice

Balb/C or C57BL/6 mice were anesthetized with a mixture of 2% Xylazine (Rompun 2% injection solution, Bayer, Germany, cat. no. QN05CM92) and 100 mg/ml Ketamine (Ketamine 10%, Parke-Davis, Germany, cat. no. FS1670044WDD). Under deep anesthesia, a ventral midline incision was performed to expose both the portal vein and the inferior vena cava. The inferior vena cava was cannulated with a 24 G catheter (Becton Dickinson, Germany, cat. no. 381312) and perfused, first with EGTA buffer at 7 ml/min, followed by collagenase II buffer (Worthington Biochemical Corporation, USA, cat. no. LS004176) at 4 ml/min for approximately 5 minutes each in healthy livers or 8 minutes each in injured livers. To relieve internal pressure, the portal vein was incised at the start of the perfusion. Following the two-step digestion procedure, the gallbladder was resected, and the digested liver was removed. Liver tissue was minced under a sterile hood in a petri dish containing 12 ml of suspension buffer and filtered through a 100 µm cell strainer (Corning, Germany, cat. no. 352360) to remove tissue debris. Hepatocytes were separated by centrifugation for 2 minutes at 50 g (hepatocytes precipitate into the pellet) and further purified using Percoll solution (GE Healthcare, Germany, cat. no. GEHE17-089-01), as described by (19). The supernatant containing non-parenchymal cells (NPC) was centrifuged twice for 2 minutes at 50 g to eliminate contaminating hepatocytes. NPC were subsequently sedimented by centrifugation for 10 minutes at 500 g, then resuspended in 1.2 ml Dulbecco’s Modified Eagle Medium (DMEM, Sigma Aldrich, Germany, cat. no. D1145) without phenol red. To enrich viable NPC, the suspension was mixed with 4.8 ml of 30% Histodenz solution (Sigma Aldrich, Germany, cat. no. D2158) and overlaid with 2 ml of phosphate-buffered saline (DPBS, Thermo Fisher Scientific, Germany, cat. no. 14190169). The mixture was centrifuged for 25 minutes at 4°C and 1500 g (no brake). After centrifugation, NPC were collected from the interphase, resuspended in 5 ml MACS buffer (Miltenyi Biotec, Germany, cat. no. 130-117-336), and passed through a 30 µm nylon mesh (Miltenyi Biotec, Germany, cat. no. 130-098-458) to generate a single-cell suspension. The suspension was centrifuged for 10 minutes at 500 g. NPC were then sequentially separated using magnetic-activated cell separation (autoMACS Pro Separator, Miltenyi Biotec, Germany, cat. no. 130-092-545), starting with the least abundant cells (HSC), followed by KC and LSEC. Briefly, the pellet was resuspended in 370 µl MACS buffer, incubated with 15 µl FcR Block and 15 µl CD271-PE antibody for 15 minutes at 4°C. The tube was inverted every 5 minutes and washed with 4 ml MACS buffer. After centrifugation at 500 g for 5 minutes, the pellet was resuspended in 180 µl MACS buffer and incubated with 20 µl magnetic anti-PE microbeads for 15 minutes at 4°C. Following additional washes, the suspension was applied to the autoMACS separator using the #POSELD2 program. CD271-positive cells (HSC) were collected, and viability and cell number were assessed using MACSQuant Analyzer 10 (described in the "Cell Yield and Viability" section). The CD271-negative cells (flowthrough) were processed similarly with CD11b microbeads for KC separation and CD146 microbeads for LSEC separation, following the protocols by (20) and (21). For Mdr2*KO* mice, livers were dissociated using the gentleMACS Dissociator (Miltenyi Biotec, Germany, cat. no. 130-096-427) with the Dissociation Kit (Miltenyi Biotec, Germany, cat. no. 130-110-201), according to the manufacturer’s recommendations. The liver tissue was processed with enzyme D, R, and A in a gentleMACS C tube (Miltenyi Biotec, Germany, cat. no. 130-093-237) using the 37°C_Multi_B_01 program for 1 hour. NPC were isolated and separated as described for Balb/C mice, using the autoMACS separator with CD271, CD11b, and CD146 markers (Supporting table 1).

### Isolation of hepatic stellate cells (standard Nycodenz-based method)

To compare the standard protocol using a Nycodenz gradient with our new CD271- based method, hepatic stellate cells (HSC) were isolated from Balb/C mice following the protocol of (11) with slight modifications. Briefly, mice were anesthetized and surgically dissected as described earlier. The liver was perfused with sterile EGTA solution for 2.5 minutes, followed by Pronase E buffer (Merck, Germany, cat. no. 1074330005) for 5 minutes and Collagenase D buffer (Roche, Germany, cat. no. 11088882001) for 8 minutes, at a pump rate of 5 ml/min. After this three-step digestion procedure and removal of the gallbladder, the liver was excised, carefully minced, and transferred to a stirring solution containing Pronase E, Collagenase D, and DNase I (Roche, Germany, cat. no. 11284932001). The solution was stirred gently at 100 rpm for 5–10 minutes. Importantly, the pH of the cell suspension was maintained at 7.4 to ensure a high yield of viable HSC.

Subsequently, the cell suspension was passed through a nylon mesh and centrifuged as previously described. The cell sediment was resuspended in 8.27% Nycodenz solution (Axis Shield, Scotland, cat. no. 1002424), overlaid with 1.5 ml of Hank’s balanced salt solution (HBSS) (Sigma Aldrich, Germany, cat. no. H6648), and centrifuged at 1400 g for 25 minutes at 4°C without brake. HSC were carefully collected from the interphase and resuspended in DMEM supplemented with 10% fetal bovine serum (FBS, Gibco, Germany, cat. no. 10270-106), 1% penicillin/streptomycin (P/S, Sigma Aldrich, Germany, cat. no. P0781), and 1% L-glutamine (Gibco, Germany, cat. no. A2916801).

### Human liver tissue and cell separation for dissociation

Healthy liver tissue from four patients who underwent hepatectomy for metastatic tumors was obtained from the local surgery department (Medical Faculty Homburg, Germany). Written informed consent was provided by all patients prior to surgery. The study protocol was approved by local ethics committees (143/21, 154/10, and 169/02). Portions of the same tissue were fixed in 4% PFA overnight at 4°C, followed by paraffin embedding for histological analysis. Liver tissue pieces were preserved in MACS tissue storage solution (Miltenyi Biotec, Germany, cat. no. 130-100-008) for approximately 18 hours at 4°C. For liver cell purification, tissue was placed in a petri dish and cut into 4 g pieces. The premixed working solution of the Tumor Dissociation Kit (Miltenyi Biotec, Germany, cat. no. 130-095-929) was prepared according to the manufacturer’s protocol where 5 ml DMEM mit 10 % Panexin, 5 % HPL, +100 µl Pen/Strep, +250 µl Enzym H+150 µl Enzym R,+ 20 µl Enzym A was mixed. Enzymatic, mechanical, and heat dissociation was performed using the gentleMACS Octo Dissociator for 65 minutes with the 37°C_h_TDK_1 program. The resulting cell suspension was filtered through a 70 µm Smart Strainer (Miltenyi Biotec, Germany, cat. no. 130-098-462), centrifuged for 5 minutes at 70 g and 4°C, and hepatocytes (HC) were collected as the pellet. The supernatant containing non-parenchymal cells (NPC) was processed as described in the mouse liver cell isolation protocol, including AutoMACS separation using human-specific magnetic bead-labeled antibodies (Supporting Table 1).

### Liver cell isolation using perfusion pipeline

Liver tissues for PHH isolation were obtained from patients undergoing hepatic resections at Leipzig University Medical Center. The study was conducted in accordance with the principles of the Declaration of Helsinki and approved by the Ethics Committee of the Medical Faculty of Leipzig University (registration number (006/17-ek, date 2017/03/21 ratified 2019/02/12). PHH and corresponding hepatic NPC were isolated from macroscopically tumor-free tissue samples by a two-step EGTA/collagenase perfusion technique as described previously (22) as summarized in supporting Table 2. In brief, after isolation, cell pellets were washed in DPBS with Ca^2+^, Mg^2+^ (Gibco, Thermo Fisher Scientific, USA, cat. no. 14040174), centrifugated at 51 xg for 5 min at 4°C to separate PHH (pellet) from NPC (supernatant). PHH were resuspended in cooled William’s Medium E (Gibco, Thermo Fisher Scientific, cat. no. 32551087), supplemented with 10% FBS (Sigma-Aldrich, USA, cat. no. S0615-500ml), 15 mM HEPES (cat. no. 15630–056), 1 mM sodium pyruvate (cat. no. 11360– 039), 1% Penicillin/Streptomycin (cat. no. 15140122), 1% MEM NEAA (cat. no. 11140– 035, all provided by Gibco, Thermo Fisher Scientific, USA), 1 µg/mL Dexamethasone (Jenapharm/MIBE GmbH, Germany, cat. no. PZN: 08704404), and 32 U/L human insulin (Eli Lilly, Germany, cat. no. PZN: 02526396). The supernatant containing the NPC was centrifuged at 300 xg for 5 min and then 650 xg for 7 min at 4°C. The resulting cell pellets were combined using HBSS (Gibco, Thermo Fisher Scientific, USA, cat. no. 14175-053). Viable PHH and NPC were counted with the trypan blue exclusion technique using a Neubauer counting chamber. 50 mio viable cells of isolated PHH or NPC were resuspended in ∼1,8 ml ChillProtec plus® medium (Merck, Germany, cat. no. F2293) and were shipped overnight (23, 24). The supernatant containing non-parenchymal cells (NPC) was processed as described in the mouse liver cell isolation protocol, including AutoMACS separation using human-specific magnetic bead-labeled antibodies (Supporting Table 1).

## Statistical analysis

Data were analyzed using GraphPad Prism version 8.0 for Windows (GraphPad Software). Results were shown as mean ± standard deviation (SD). Two-tailed unpaired Student t-test or one-way ANOVA was used to compare between two groups or more groups. Probability value (p) less than 0.05 was considered significant and represented graphically as *p<0.05; **p<0.01; ***p<0.001, unless indicated otherwise. Additional data and methodology can be found in the Supporting material section.

## Results

### Confirmation of CD271 as a HSC specific surface marker in healthy and diseased mouse and human livers

CD271 has been identified as HSC surface protein (16). To investigate localization of CD271 expression in both mouse and human livers under healthy and diseased conditions, we examined its expression with other HSC markers in liver tissue. In chronically diseased mouse livers (CCl_4_ and Mdr2*KO* mice), we observed that a co-localization of CD271 and desmin in both healthy and diseased livers (Figure 1A and B, upper panel yellow arrows; Supporting figure 1). However, this co-localization was more prominent in areas of tissue damage, where CD271 also co- localized with α-SMA (Acta2) expression (Figure 1A and B, lower panel). To further support these findings, we also analysed the publically available datasets from CCl_4_-treated (Figure 1C) and Mdr2*KO* (Figure 1D) mouse liver arrays. These datasets confirmed that CD271 RNA expression closely parallels that of desmin, reinforcing the notion that CD271 is a marker of both activated and quiescent HSC, in line with the co- localization results in tissue sections. Similar observations were made in human liver tissue, where CD271, desmin and ACTA2 co-localized in both healthy and fibrotic regions (Figure 1E; Supporting Figure 2). Analysis of array datasets also showed similar patterns of CD271 expression (Figure 1F) in different fibrosis stages (indicated by Scheuer scoring). Additionally, serial sections of human healthy livers were subjected to IHC staining with anti-CD271 and anti-ACTA2 antibodies, as well as IF staining for desmin. Co-localization of CD271 and ACTA2 was observed in perisinusoidal and fibroblast-like cells (Supporting Figure 3), and in close proximity to desmin-positive cells, further supporting the role of CD271 as a marker of activated HSC. In conclusion, CD271 is confirmed as HSC surface marker in both healthy and diseased mouse and human livers, where it co-localizes with well-known HSC markers like desmin and ACTA2.

**Figure 1:**
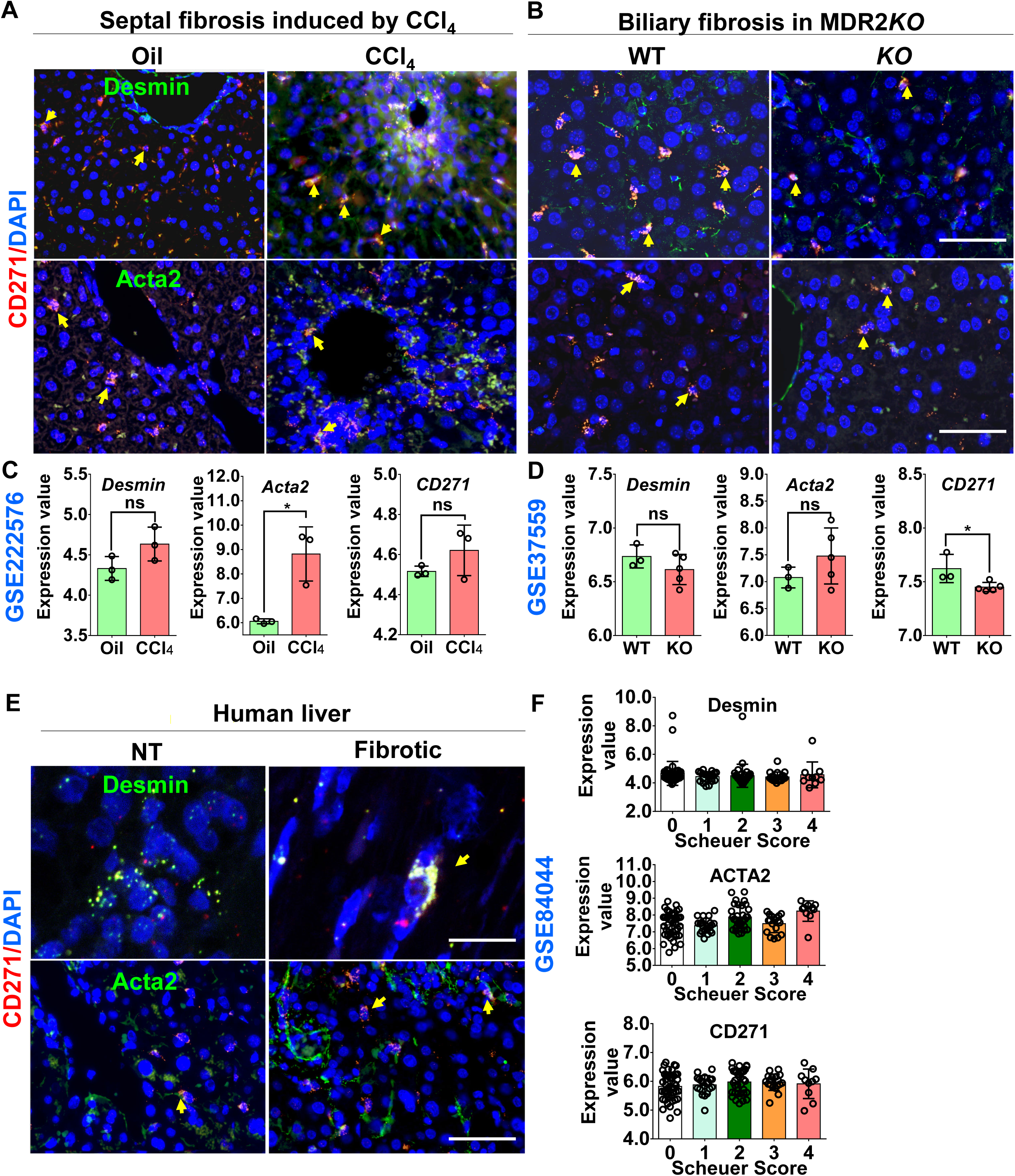
CD271 is a specific marker for quiescent and activated HSC. (A, B) Immunofluorescence (IF) staining using antibodies against desmin (a marker for all HSC) and Acta2 (a marker for activated HSC) coupled with RNAscope analysis for CD271 in mouse models of liver fibrosis. Septal fibrosis was induced using repeated intoxication with CCl_4_ treatment, and biliary fibrosis was modeled in Mdr2*KO* mice. Yellow arrows indicate co-localization of the positive signals. Scale bars are 100µm. (C, D) Transcriptomic analysis of CD271, Acta2, and desmin expression in CCl_4_-induced fibrosis and Mdr2*KO* livers using publicly available datasets (GSE222576 and GSE37559, respectively). (E) Tumor-free (NT) and fibrotic human liver samples were co-stained with RNAscope for CD271 and antibodies against Acta2 or RNAscope for CD271 and desmin. Colocalization is marked by yellow arrows. Scale bars are 100µm. (F) additionally, publicly available data (GSE84044) were analyzed to compare CD271, Acta2, and desmin expression with the Scheuer fibrosis score in human livers. ns: not significant; *p<0.05 compared with controls.

**Figure 2:**
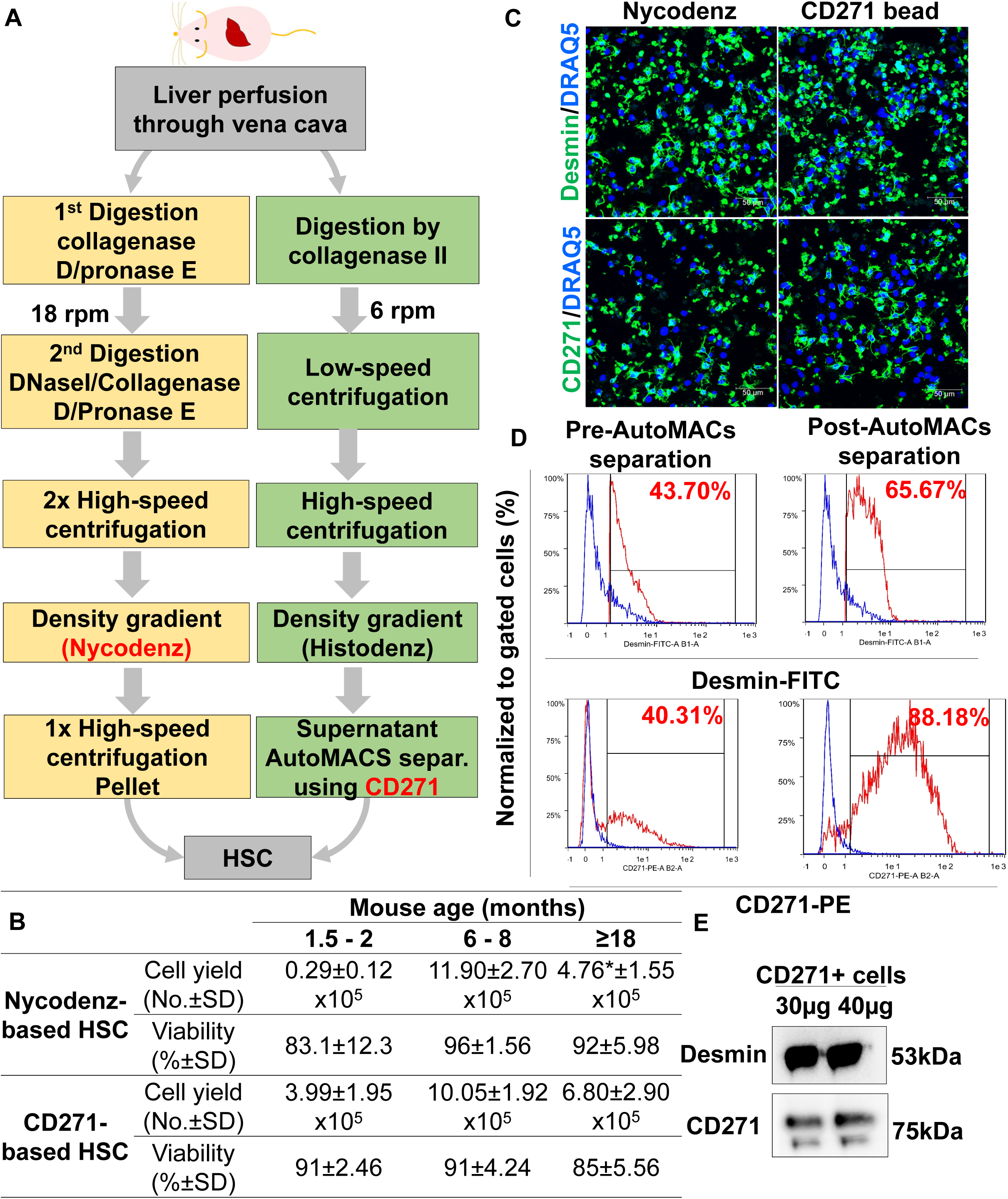
**Comparative HSC separation with the classical Nycodenz-based method and our new CD271-labelled magnetic bead-based method. (**A) Flow chart of the different HSC isolation procedures from Balb/C. Livers were perfused through the vena cava inferior and digested by collagenase D, DNase and pronase E in the “Nycodenz method” (left panel), whereas only collagenase II was used in the CD271 based procedure (right panel). rpm: round per minute (pump speed). (B) Comparative analyses of HSC yield, viability and purity with the Nycodenz or CD271 sorting method from livers of Balb/C mice at different ages, as indicated. (C) Immunofluorescence staining shows desmin staining in CD271-positive cells, as is the case for Nycodenz- purified cells, in both cases from 6-8 months old Balb/C mice. Scale bars represent 50μm. (D) MACS flow cytometry analysis using desmin and CD271 markers before and after CD271-bead AutoMACs separation indicating enrichment of HSC. Percentages and red histogram indicate the desmin^+^ or CD271^+^cells. Blue histogram indicates autofluorescence cells. (E) Immunoblot analysis of CD271 and desmin of cells isolated by CD271 beads AutoMACs separation indicates their co-expression in HSC. The numbers are means ±SD of 4-6 mice per condition, and representative images are shown. *Not detected due to insufficient yield in young mice or contamination with lipid droplet-containing hepatocytes in aged mice, respectively.

**Figure 3:**
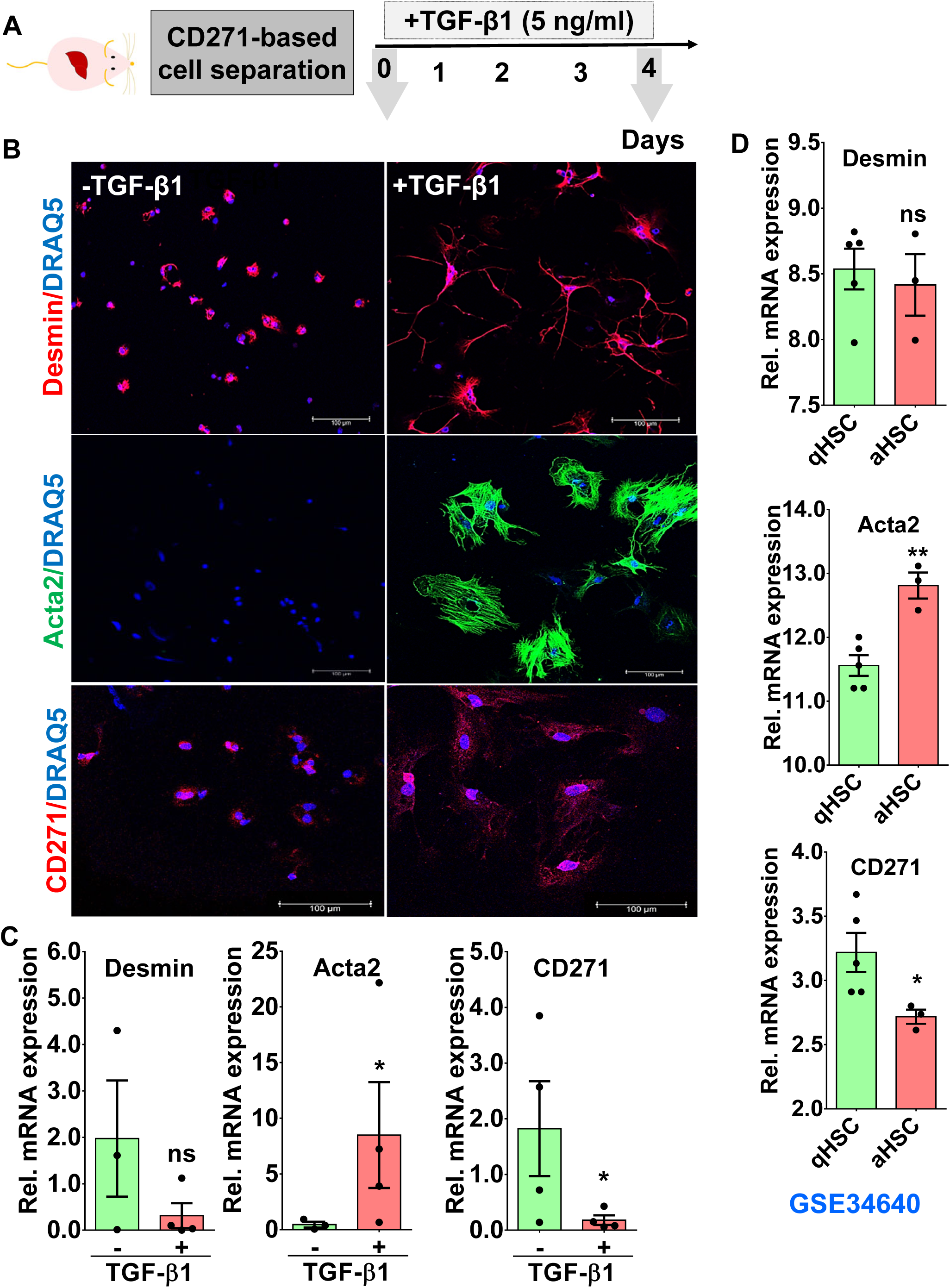
**CD271-separated cells respond to TGF-β1 as functional HSC**. (A) CD271-separtated cells were incubated with TGF-β1 for 4 days. (B) Immunofluorescence staining of CD271-separated cells upon TGF-β1 stimulation for 4 days. Desmin and Acta2 as markers for total and activated HSC, respectively, are observed in CD271-separated cells. CD271 is also observed in both HSC phenotypes. Scale bars represent 100μm. (C) RT-PCR analysis of *Desmin, Acta2* and *Cd271* cells separated by CD271-beads upon TGF-β1 stimulation. Representative images and numbers from n=3-6 animals per condition of three independent experiments. (D) Transcriptomics analysis of publically available dataset (GSE34640; (25) of *Desmin, Acta2* and *Cd271* expression. In this report, HSC were isolated using Nycodenz density centrifugation and stimulated in plastic dishes using FBS for 64h. aHSC: activated hepatic stellate cell; qHSC: quiescent hepatic stellate cell. ns: not significant; *p<0.05; **p<0.01 compared with controls.

### Confirmation of CD271 as strain and age-independent specific HSC surface marker

We first aimed to test specificity and reliability of CD271 for isolating pure HSC from mouse liver. Conventional Nycodenz- (according to (11)) and the new CD271- magnetic bead-based HSC isolation methods were compared by using healthy Balb/C mice at different ages (isolation schemes shown in Figure 2A). Cell viability was throughout higher than 85% in all mice for CD271-based isolation (Figure 2B). The purified HSC were seeded and subsequently examined for desmin positivity (as conventional HSC marker) via IF staining (Figure 2C). With both protocols, most (>88%) cells were desmin-positive, indicating high purity. Noteworthy, conventional and CD271-based methods were similarly effective in middle-aged mice (6-8 months old) with a yield of about 11.90±2.70×10^5^ and 10.05±1.92×10^5^ HSC per mouse, respectively (Figure 2B). However, the CD271 based purification method was superior in young mice (1.5-2 months) producing 13 times more HSC, where the HSC yield was about 3.99 ±1.95×10^5^ per mouse as compared to only 0.29±0.12×10^5^ HSC with the Nycodenz method (Figure 2B), the latter not being sufficient in numbers for most downstream analyses, e.g. for biological assays. Furthermore, the CD271 method yielded a 40% higher HSC number in aged mice (1.5 years old) compared to the Nycodenz method (Figure 2B), which failed due to contamination with fat-laden hepatocytes. It is generally accepted that C57Bl/6 mice are not well suited for HSC isolation due to insufficient yield when using Nycodenz-based method. This phenomenon is a major drawback in liver research, since C57Bl/6 mice are widely used for *in vivo* studies. To gain knowledge about the degree of enrichment upon cell separation using CD271-AutoMACs technology, we used MACS flow cytometry to determine the enrichment of HSC fraction. After exclusion of dead cells by propidium iodide (PI) positive cells (Supporting Figure 4A), debris (Supporting Figure 4B), and unstained cells were gated by high levels of APC and APC-Vio777 to exclude autofluorescent cells (Supporting Figure 4B-C). The remaining cells were simultaneously labelled with desmin-FITC and CD271-PE antibodies and analysed by MACs flow cytometry. We found two populations of separated cells based on desmin^high^/CD271^high^ (Figure 2D; red histogram) and desmin ^low^/CD271^low^ (Figure 2D; blue histogram). We found that the CD271-magnetic bead autoMACs separation further enriched desmin^high^/CD271^high^ cells approximately about 1.5 and 2.2 fold, respectively, compared to pre-separated cells (Figure 2D; red: post-autoMACS separation). Thus, autoMACs separation based on CD271 labelling reliably separates and enriches the desmin positive HSC fraction. In addition, immunoblot analysis shows that both markers can be detected in the HSC fraction (Figure 2E). In conclusion, the CD271-based isolation method proves to be a superior and versatile tool for efficiently isolating pure HSC from mouse livers, with strong applicability across different age groups and mouse strains, offering enhanced purity and yield compared to traditional methods.

**Figure 4:**
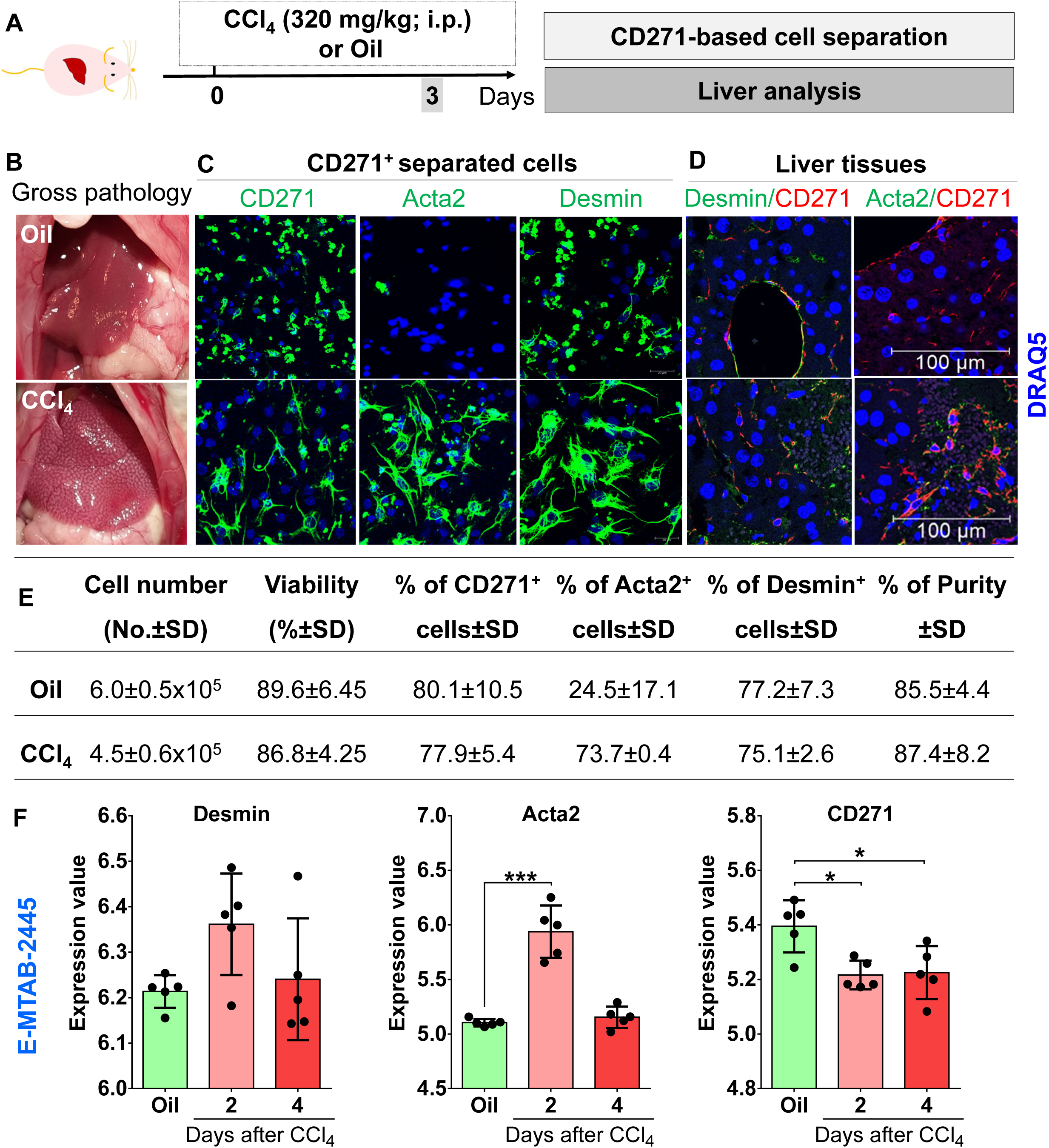
**CD271-separated cells are quiescent and activated HSC. (**A) Experimental scheme. (B) Gross pathology of livers at day 3 after oil or CCl_4_-exposure of BalbC mice. (C) Immunofluorescence staining using CD271 and desmin or Acta2 of CD271- isolated cells from oil and CCl_4_-exposed livers. Scale bars are 20μm. (D) Co- immunofluorescence staining using CD271 and desmin or Acta2 of liver tissues. Scale bars are 100μm. (E) Comparative analyses of HSC yield, viability and purity of CD271 sorting method from livers of Balb/C mice, as indicated. (F) Transcriptomics analysis of publically available dataset (E-MTAB-2445; (28) of *Desmin, Acta2* and *Cd271* expression. In this report, mRNA levels of *Desmin,* Acta2, and *Cd271* is analysed upon oil or CCl_4_ exposure at day 2 and 4. Representative images are from n =3-4 animals per condition of three independent experiments. ns: not significant; *p<0.05; ***p<0.001 compared with controls.

### CD271 separated cells show characteristic HSC behavior

To functionally prove the characteristic HSC phenotype of isolated CD271 positive cells, a TGF-β1 treatment experiment was conducted (Figure 3A). As TGF-β is a potent activator of HSC, the isolated cells were expected to transdifferentiate into myofibroblasts. By morphology, TGF-β1 treated cells acquired a fibroblast phenotype after daily treatment for 4 days (Figure 3B). Desmin, a marker for quiescent and activated cells, was repressed whereas ACTA2 as an indicator of activated HSC was increased upon TGF-β stimulation (Figure 3C). Noteworthy, Cd271, similar to Desmin, was downregulated in stimulated HSC (Figure 3C). This is in an agreement with publically available transcriptomics datasets (25) from HSC which were isolated by Nycodenz density centrifugation. In these datasets, mRNA levels of Desmin and Cd271 were repressed upon activation of HSC via culture on stiff plastic or by co-culture with KC (Figure 3D). These activation processes, similar to TGF-β effects, were accompanied by induction of ACTA2 expression (Figure 3D). Hence, the CD271-magnetic bead semi-automated method is a reliable method to separate functional, TGF-β-responsive, HSC in high yield and purity.

### CD271 is a specific marker for HSC separation from diseased livers

During the process of activation, HSC lose fat droplets and become fibroblast-like cells. To evaluate the effectiveness of the CD271-bead method for isolating activated HSC, we isolated cells 72h after CCl_4_-induced acute liver injury (Figure 4A) in Balb/C mice. As expected, necrotic foci were observed in the CCl_4_-treated tissues during gross pathology analysis (Figure 4B). Next, the CD271-based cell separation was performed. Immunofluorescence staining revealed that CD271+ cells exhibited high desmin positivity in both oil- and CCl_4_-treated livers (Figure 4C). Acta2 positivity of CD271^+^ cells were exclusively observed in CCl_4_-exposed livers (Figure 4C). This finding indicates that the separated cells were desmin-positive in both healthy and CCl_4_- exposed livers, with Acta2 expression restricted to cells from CCl_4_-treated livers (Figure 4C). Co-immunofluorescence showed a strong co-localization of CD271 with desmin was found in both oil (healthy) and CCl_4_ exposed mice (Figure 4D). Cell viability exceeded 86% across all mice. HSC yields were 6±0.5×10⁵ and 4.5 ± 0.6×10⁵ cells per liver in oil- and CCl_4_-exposed groups, respectively (Figure 4E). The purity of CD271- separated cells was consistently above 85% (Figure 4E). To confirm CD271 expression in CCl_4_-exposed mouse livers, publicly available datasets were analyzed, revealing that CD271 displayed a similar expression pattern to other HSC markers at days 2 and 4 post-injection (Figure 4F). These findings demonstrate that CD271 is a reliable marker for isolating hepatic stellate cells from both healthy and diseased livers. The high viability, yield, and purity of CD271^+^ cells further highlight the effectiveness of this method.

### Simultaneous isolation of different liver cell types from healthy and diseased mouse livers

After demonstrating that CD271 is a reliable marker for isolating HSC from both healthy and diseased murine livers, the next step was to develop a semi-automated pipeline using magnetic bead-labelled antibodies to purify hepatocyte and other non-parenchymal cell (NPC) populations from a single mouse liver. For this, we used 6-8-month-old Balb/C mice and combined manual processing from previous protocols (11) with magnetic bead-activated cell separation (MACS) into a single workflow. A schematic of the process is shown in Figure 5A. Following the initial low-speed centrifugation step for hepatocytes separation, NPCs were purified using magnetic bead-labelled antibodies targeting CD271 (HSC), CD11b (KC), and CD146 (LSEC), as depicted in Figure 5A.Cell yields for hepatocytes, HSC, KC, and LSEC were 33.4 ± 5.5×10^6^, 51.2 ± 6.3×10^4^, 12.4 ± 4.8 ×10^5^, and 18.2 ± 8.9×10^5^ cells per healthy mouse liver, respectively (average ± SD of 4-6 mice; Figure 5B). Cell viability, assessed by trypan blue staining, was 92.2 ± 2.8% for hepatocytes, 86.6 ± 5.9% for HSC, 91.1 ± 5.1% for KC, and 89.3 ± 4.6% for LSEC (Figure 5B). To assess purity, immunofluorescence staining for albumin (hepatocyte), desmin (HSC), F4/80 (KC), and Lyve1 (LSEC) was performed (Figure 5C). The isolated cells showed high purity: 98.6 ± 0.5% for hepatocyte, 88.9 ± 7.8% for HSC, 87.0 ± 10.4% for KC, and 89.1 ± 11.4% for LSEC (Figure 5B and C). We further evaluated CD271 as a specific surface marker for HSC by immunoblotting, confirming its expression exclusively in the desmin-positive cell population (Figure 5D). Additionally, CD31 and F4/80, well- established markers for LSEC and KC, respectively, validated the purity of the KC and LSEC fractions (Figure 5D). The purity of isolated LSEC is evaluated by LYVE1 (specifically expressed in LSEC, not in the endothelial cell of large vessels) as suggested by (20). At the mRNA level, desmin and Cd271 expression were highest in the HSC fraction compared to hepatocytes, KC, and LSEC (Figure 5E). Further RT- PCR analysis of albumin (hepatocyte marker), CD11b (KC marker), and Lyve1 (LSEC marker) (Figure 5E) confirmed the exclusive expression of these markers in their respective cell types, further validating the high purity of the isolated populations. Cell isolation success is critically dependent on mouse strain and age. To evaluate the versatility of our pipeline, we applied it to 6-8-month-old C57Bl/6 mice (Supporting Figure 5A). In addition to the efficient isolation of CD271^+^ HSC, the protocol also enabled the effective purification of hepatocyte, LSEC, and KC (Supporting Figure 5A and 5B). We further tested the isolation procedure on young adult (2-3 months) and aged (18 months) mice. Cell yield data showed that the protocol is applicable to these age groups as well, producing pure cell fractions with high yields and viability, suitable for subsequent analysis (Supporting Figure 5B). To further assess the robustness of our isolation pipeline in diseased models, we applied it to the Mdr2*KO* model of chronic biliary liver disease at 4 months of age (Supporting Figure 5C). The magnetic bead- labelled isolation method yielded satisfactory cell numbers for HSC, LSEC, and KC, with approximately 90% viability (Supporting Figure 5D). Compared to healthy Balb/C mice, Mdr2*KO* mice showed reduced HSC yields, with only 1.0 ± 0.20×10^5^ cells per mouse. To assess purity, mRNA expression of cell-type markers was analyzed. As expected, albumin was exclusively expressed in the hepatocyte fraction, CD271 was restricted to the HSC population, CD11b was exclusively found in the KC fraction, and Lyve1 expression was significantly higher in the LSEC, confirming the efficacy of the magnetic bead-based isolation procedure in diseased livers (Supporting Figure 5E). These results demonstrate that our semi-automated liver cell isolation pipeline is not only efficient for isolating major liver cell types with high yield and excellent purity from healthy mouse livers, but is also applicable to diseased models, making it a valuable tool for studying liver pathology.

**Figure 5:**
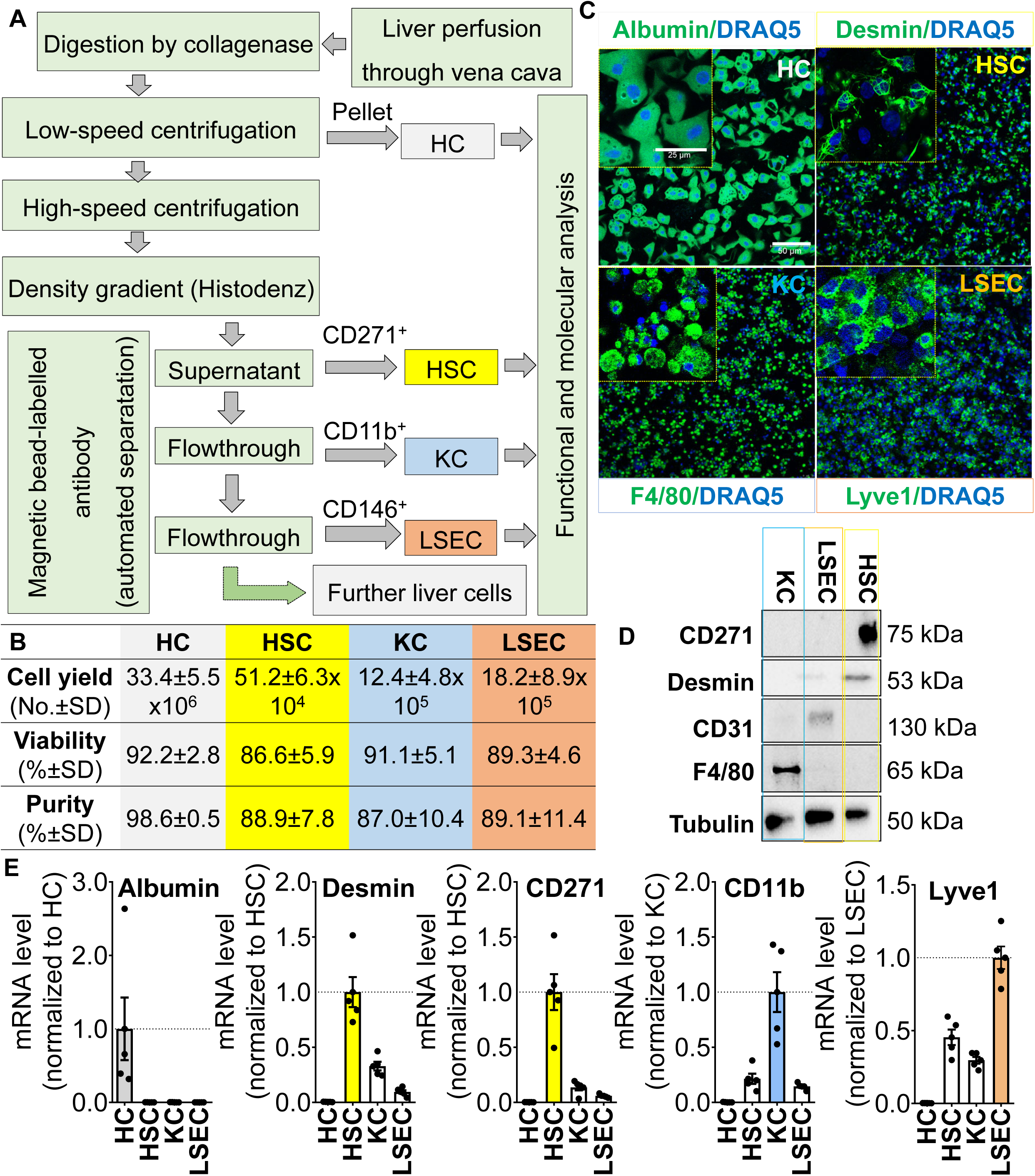
Isolation of the four liver cell populations from a healthy mouse liver.. (A) Proposed pipelines for simultaneous isolation of different cell types from one mouse liver. Balb/C mouse liver is perfused through the vena cava inferior and digested by collagenase. After centrifugation, the pellet contains hepatocytes (HC) and supernatants are subjected to magnetic beads-labelled antibodies namely CD271, CD146 and CD11b for automatic separation of hepatic stellate cells (HSC), Kupffer cells (KC) and liver sinusoidal endothelial cells (LSEC), respectively. (B) Number, viability and purity (based on quantification of IF staining of respected markers) of isolated cells by semi-automated pipelines from one healthy mouse liver. (C) Immunofluorescence staining of cell specific markers including albumin (HC), desmin (HSC), Lyve1 (LSEC) and F4/80 (KC). Scale bars are 50μm and 25μm for overviews and closeups, respectively. (D) Immunoblot analysis of CD271, desmin, CD31 and F4/80 in KC, LSEC and HSC. (E) mRNA levels of *Albumin, Desmin, CD271, Lyve1* and *Cd11b*. Representative images and numbers from n =4-6 animals per condition of three independent experiments.

### Isolation of four liver cell types from human liver

We applied the magnetic bead-based MACS separation system to purify liver cells from freshly resected human livers. Human liver tissue was either processed immediately using a conventional perfusion-based method as described by (24) or stored in MACS storage buffer upon resection and subsequently processed for dissociation (workflow illustrated in Figure 6A). Using the conventional perfusion-based method, liver tissue was perfused, and after low speed centrifugation hepatocyte fraction underwent Percoll purification to discard dead cells. The supernatant was subjected to high-speed centrifugation and followed by magnetic bead separation to isolate HSC, KC, and LSEC. The yields achieved were 14.2±6.6×10^6^ hepatocytes, 1.7±0.1×10⁵ HSC, 4.9±1.5×10⁵ KC and 2.5±0.7×10⁵ LSEC per liver piece, with cell viability exceeding 80% for all NPC fractions (Figure 6B). Purity of the isolated cell populations was confirmed at the mRNA level using specific markers: albumin (hepatocyte), desmin (HSC), CD271 (HSC), CD11b (KC), and CD146 (LSEC) (Figure 6C). In contrast, the dissociation-based method showed suboptimal purity (Figure 6D) for non-parenchymal cells (NPCs).

**Figure 6:**
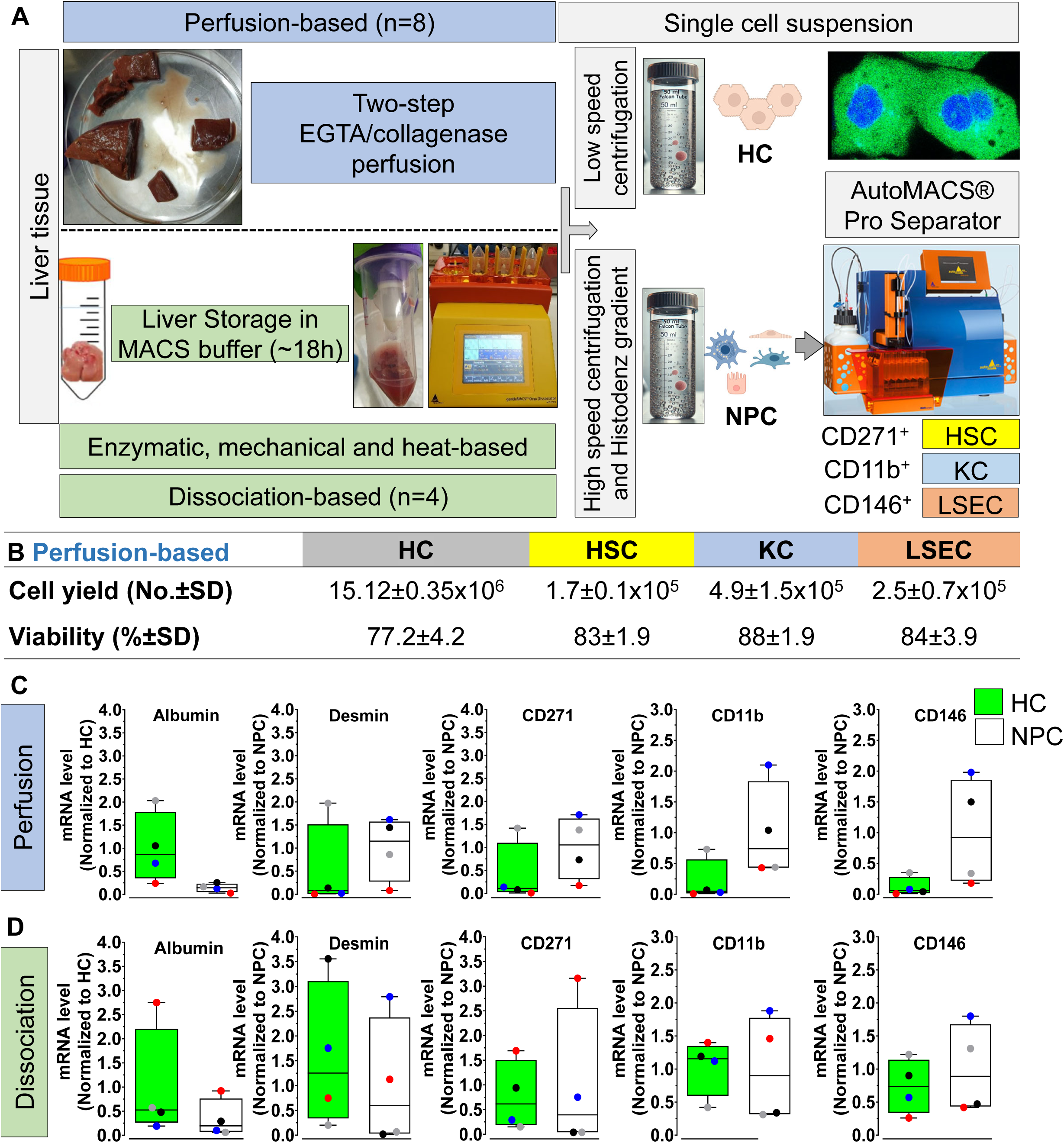
Isolation and characterization of human livers cells. (A) Workflow illustrating the steps for isolating hepatocytes (HC), and non-parenchymal cells (NPC) including hepatic stellate cells (HSC), Kupffer cells (KC), and liver sinusoidal endothelial cells (LSEC) from liver tissue using two different methods for single cell preparation. Perfusion-based method requires a big piece of liver (>10g; n=7) and perfused with a two-step EGTA/collagenase perfusion according to (24). In case of dissociation-based method, small liver pieces (>4g; n=4) is kept in MACS tissue storage buffer (∼18h) for Enzymatic, mechanical, and heat-based dissociation. A single cell suspension for both method was similar as follows; (i) low speed centrifugation and percoll purification to isolate HC and (ii) the supernatant was subjected to high speed centrifugation and Histodenz gradient to obtain a single cell from NPC fraction. Separation using the AutoMACS Pro Separator was performed using a magnetic-bead labelled antibodies specific for HSC, KC and LSEC. (B) Yields and viability of isolated liver cell populations. Cell yields (number per gram of tissue, mean ± SD) and viability (%±SD) for HC, HSC, KC, and LSEC are shown for perfusion-based method, since dissociation-based was not optimal to separate and purify specific cell). The HC number and viability as summarized in supporting table 2. (C and D) Representative mRNA expression levels of cell-specific markers based on perfusion and dissociation- based method. Normalized mRNA levels for albumin (HC), desmin (HSC), CD271 (HSC), CD11b (KC), and CD146 (LSEC) confirm the purity of the isolated populations.

## Discussion

The liver is a multicellular organ comprising hepatocytes, hepatic stellate cells (HSC), liver sinusoidal endothelial cells (LSEC), Kupffer cells (KC), and other cells, such as cholangiocytes. Primary liver cell cultures replicate several (patho)physiological features of *in vivo* systems and offer an alternative to reduce the use of experimental animals (26, 27). Isolated cells have considerably expanded our knowledge of the (patho)biology of hepatocytes and HSC (28, 29). Several attempts have been made to isolate and characterize single liver cell populations, i.e. hepatocytes (28, 30), HSC (11, 31–33), LSEC (20, 21) and KC (34, 35). Further, protocols for simultaneous isolation of main liver cells have been developed (10, 24, 29). However, distinct disadvantages and problems occurred to achieve sufficient pure and healthy cells from small piece of liver tissue. We here describe and characterize a semi-automated protocol for the isolation of key liver cell populations, namely hepatocytes, HSC, KC and LSEC from a single mouse liver or small piece (approximately 2g) of human liver specimen. This protocol is characterized by high flexibility and adaptability. One advantage of this protocol is its applicability for mice of different ages or strain. Even more, the protocol can be applied to acutely damaged (by intoxication) and fibrotic livers (Mdr2*KO* mice) as well as human tissue. Furthermore, the protocol is extendable to include additional liver cell types, such as bile duct cells, by selecting specific surface markers. Importantly, this approach ensures high yields of pure and healthy cells, making it a versatile tool for liver cell research. HSC comprise approximately 5–8% of the total cell population in a normal liver and reside in the perisinusoidal space (Space of Disse) between hepatocytes and LSEC (1, 2). In healthy livers, HSC function as lipocytes, storing vitamin A and regulating sinusoidal blood flow (reviewed by (36)). However, during liver disease, quiescent HSC become activated, lose their lipid droplets, express Acta2, and secrete extracellular matrix proteins, contributing to fibrosis development (37). HSC are highly sensitive to stress induced by isolation procedures or culture conditions, making it challenging to develop protocols that reliably isolate HSC with other liver cell populations simultaneously. The first successful HSC isolation was conducted in rats using a combination of enzymatic digestion and density gradient centrifugation, achieving a purity exceeding 70% (38). Since then, efforts have been made to enhance HSC purity. For instance, Mederacke and co-workers employed a retrograde Pronase-Collagenase digestion of the liver followed by density-gradient centrifugation to isolate quiescent and activated HSC from both normal and fibrotic mouse livers (11). However, this Nycodenz-based protocol is optimized for Balb/c mice aged 5–7 months, as sufficient lipid accumulation in HSC is critical for effective cell enrichment via density gradient centrifugation. Additionally, HSC yield varies significantly by mouse strain; Balb/c mice produce several-fold higher HSC numbers compared to C57BL/6 mice under identical conditions (11). CD271- based isolation of HSC yields comparable numbers from different mouse strains, making it particularly suitable for isolating HSC from C57BL/6 mice—a commonly used strain for genetic studies. The Mederacke protocol, involving long digestion steps with pronase and collagenase (11), negatively impacts the viability of other liver cell types, such as hepatocytes, LSEC and KC, thereby limiting its broader applicability. In contrast, the gentle digestion with collagenase II in our protocol allows for the simultaneous isolation of HSC along with other liver cell types. Interestingly, CD271- based isolation fails when liver homogenates are prepared using Pronase and Collagenase D, likely due to degradation of the CD271 surface marker (data not shown). An alternative approach involves isolating HSC based on their autofluorescence (due to vitamin A storage), as demonstrated in rats (39) and mice (40) using FACS sorting equipped with UV lasers. However, this method is influenced by various factors, including animal diet, age, species, and genetic background, which can reduce the yield and purity of isolated cells. Furthermore, activated HSC lose lipid droplets and retinoid content, complicating isolation via autofluorescence-based methods. While FACS sorting combined with density-gradient centrifugation can achieve ultrapure HSC (33), it is costly, time-consuming, and significantly reduces cell yields. In our protocol, we tested three HSC surface markers—PDGFR-β, GFAP, and CD271—for their suitability in isolating functional HSC from healthy mouse livers using AutoMACS technology (Data not shown). CD271 emerged as the most effective marker, providing good yields of pure HSC that could also be integrated into protocols for isolating other liver cell types. CD271, a member of the tumor necrosis factor receptor superfamily, is expressed in quiescent and activated HSC in mice, humans (41) and rats (14). Its expression on HSC also support the hypothesis about their mesenchymal origin, a topic of ongoing debate (42, 43). Unlike density-gradient protocols optimized for middle-aged Balb/c mice (11), our CD271-based protocol is strain-, age-, species-, and disease-state-independent. It performs well with young or very old mice and remains effective under fibrotic conditions. Although TGF-β downregulates CD271 expression in cultured primary HSC, residual expression allows for the isolation of both quiescent and activated HSC (14, 41). We also refined the liver perfusion process, adopting a retrograde perfusion system through the inferior vena cava instead of the portal vein, simplifying cannulation and improving perfusion efficiency (11). For fibrotic mouse (based on Mdr2KO) the GentleMACS tissue dissociator, was combined with a specialized perfuser tube and liver dissociation kit, successfully isolating hepatocyte, HSC, LSEC, and KC. For healthy human livers, the GentleMACS tissue dissociator (without perfusion), combined with a specialized liver dissociation kit was not optimal to separate PC from NPC cells due to mechanical disruption, which likely damaged surface epitopes critical for effective magnetic bead labelling and separation. However, mouse hepatocyte yields were compromised, likely due to their larger size compared to human hepatocytes. Unfortunately, the dissociation-based method using 2g pieces of human tissue was suboptimal for separating liver cells, contrary to the findings of Zhai et al. (44), who suggested that HSCs could be purified through dissociation. Ultimately, this combination of mechanical and enzymatic digestion, coupled with density gradient centrifugation, produces sufficient quantities of liver cells from one mouse liver for downstream analyses. Based on the suboptimal dissociation method results, we tested a conventional perfusion-based approach (using >10 g of tissue) combined with MACS separation, confirming preserved surface marker expression and simultaneous NPC isolation. We present a reproducible, semi-automated protocol that enables the simultaneous isolation of hepatocytes, HSC, LSEC, and KC from a single mouse liver or human liver specimen based on perfusion and MACS separation. This method provides high yields of functional, pure, and viable cells, suitable for downstream applications. Although the initial setup requires significant investment in specialized equipment, the protocol’s flexibility and reproducibility make it a valuable tool for laboratories engaged in primary liver cell research. Its scalability and adaptability further expand its utility to human tissue studies based on perfusion and MACS pipeline to isolate rare liver cell types, positioning it as a versatile pipeline for advancing liver cell biology research. By leveraging the presence and preservation of surface markers, the pipeline can be further optimized to isolate rare cells or specific liver cell subtypes. Despite these advantages, the limited access to human liver tissue and significant variability, which could affect reproducibility and need further optimization. The scalability of the protocol for use in human livers or larger animals, where sample sizes are often limited or more variable, needs to be further evaluated and established.

## Supporting information

Graphical Abstract

## Acknowledgements

The authors thank Stefanie Uhlig, Vanessa Nalewaja, Christof Dormann (Medical Faculty Mannheim-University of Heidelberg, Germany) and Dr. Stefan Schnell (Flow Cytometry Application Specialist, Miltenyi Biotec, Germany) for their excellent technical assistance in FACS analysis.

## Authors contributions

AD, BD, KG and SH; experimental design and major experimental performance, data acquisition and analysis. SA, VH, PE, KG, RB, TC, CM, AD and SH; performed RT-PCR, IHC staining and imaging, assays, and analyzed the data. LK, BK, CR, MG, GD, AS, DS, EB, and NR contributed to the provisioning of human liver specimens and primary cells. AD, BD, ME, GD, CR, SD and SH; manuscript drafted, critical revised, and finalized manuscript. All authors read and commented on the final version of the manuscript.

