## supporting fig for "CD271 sorting for improved liver cell isolation: Semiautomated and simultaneous preparation of parenchymal and non-parenchymal cells from mouse and human livers"

**Supporting Fig 1**

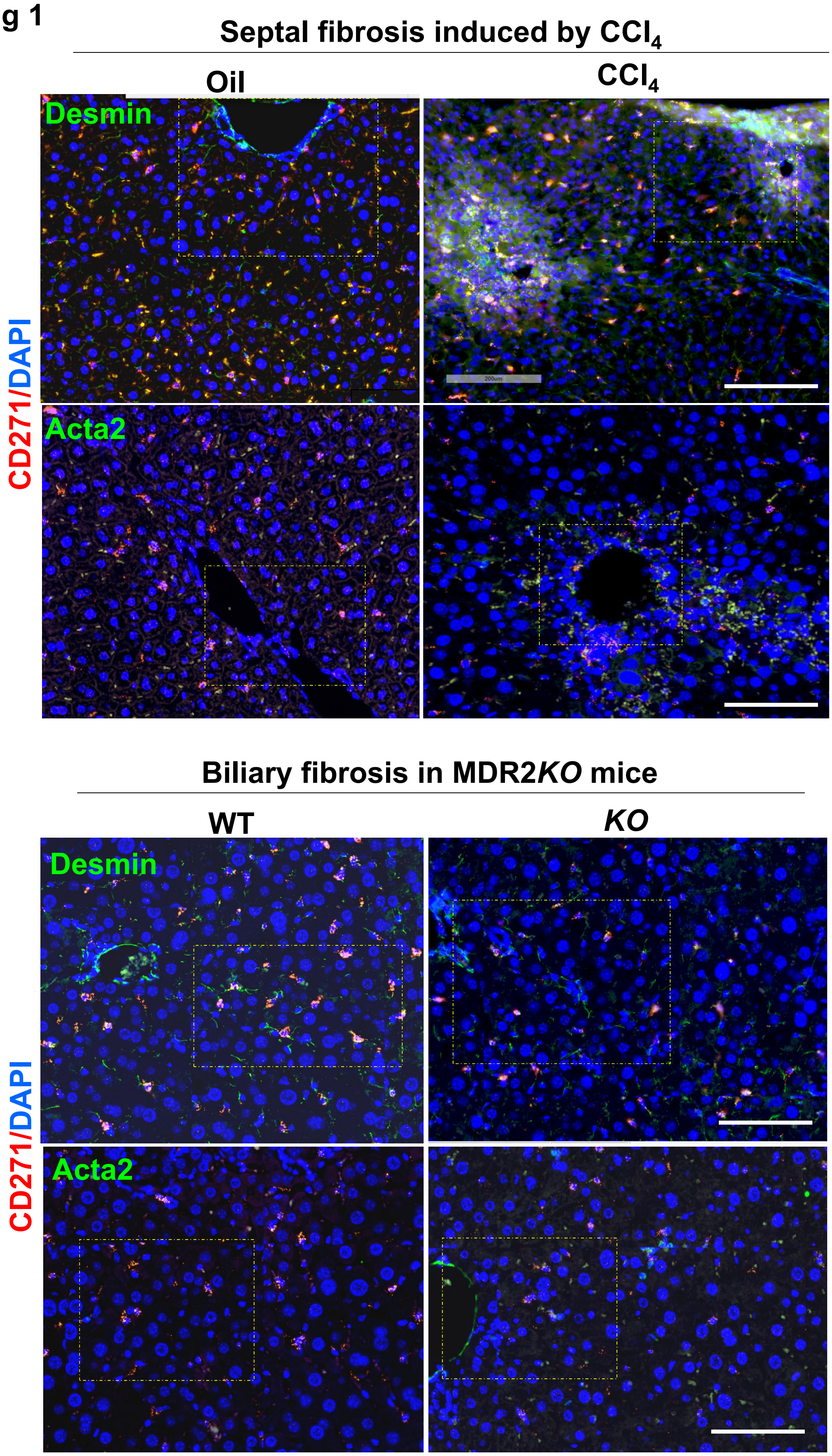

### Supporting Fig 2

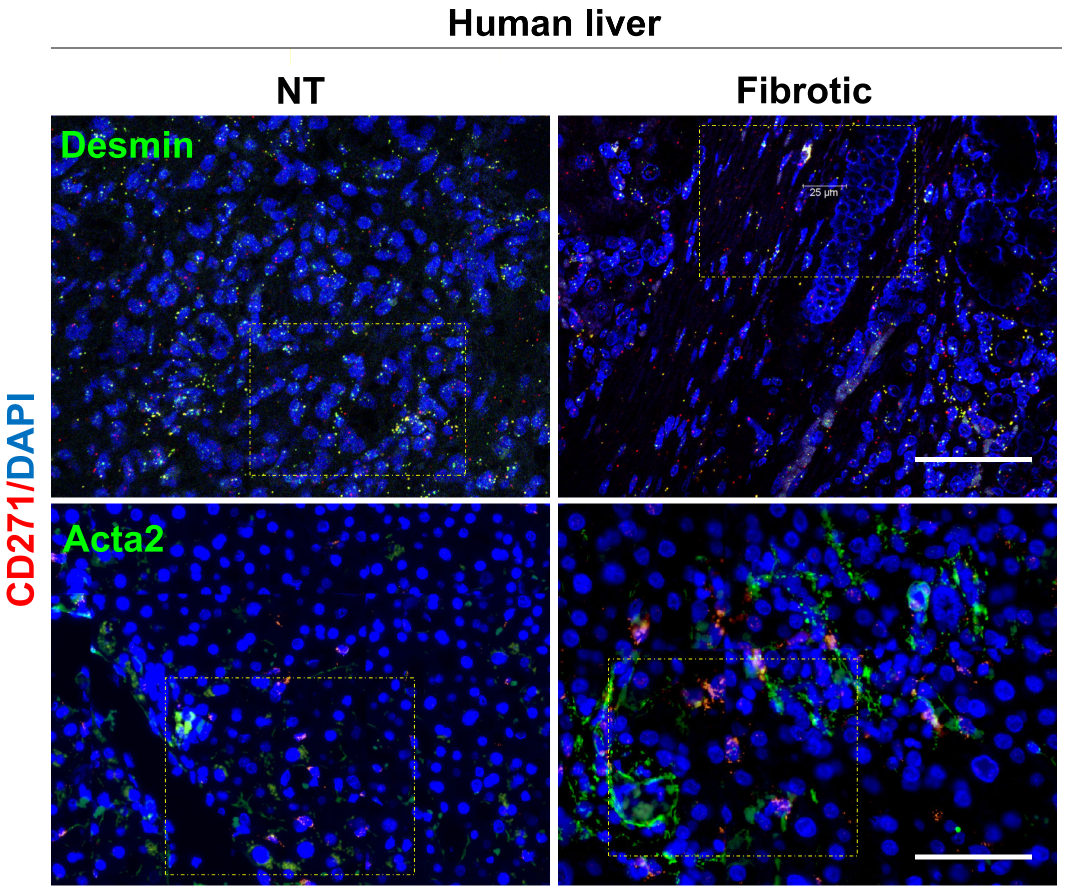

**Supporting Fig 3**

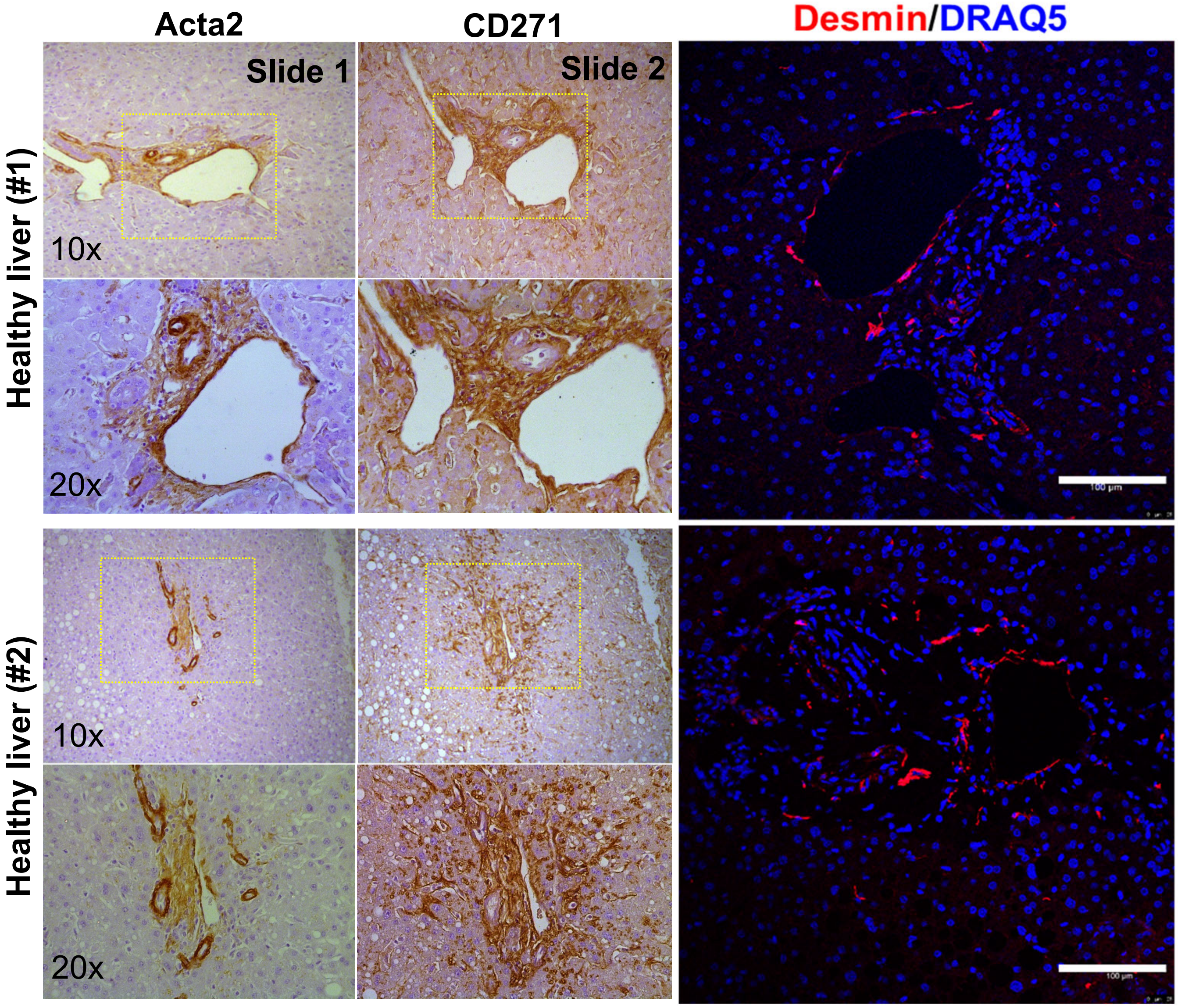

Supporting Figure 4

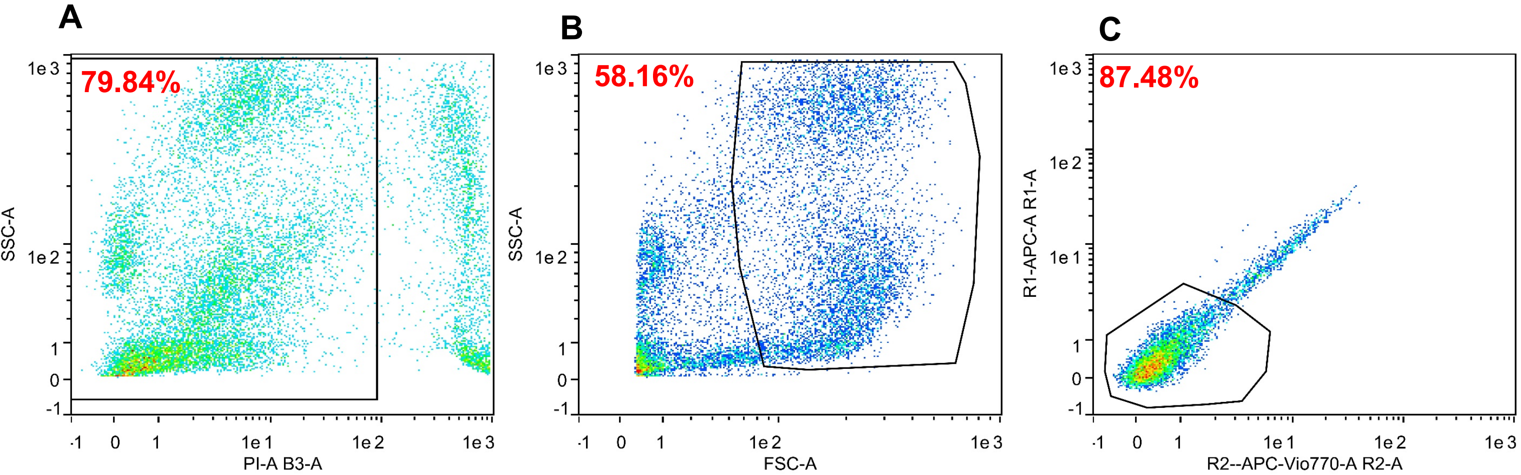

**A**

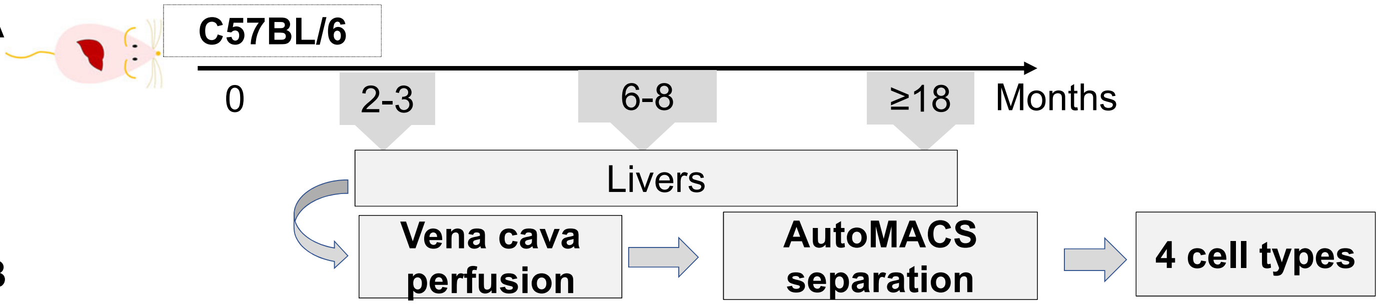


**B**

| Age (m) | | HC | HSC | KC | LSEC |
|---|---|---|---|---|---|
| 2-3 | Cell yield (No.±SD) | 35.8±2.1x $10^6$ | 122.5±44x $10^3$ | 63.8±19.6x $10^4$ | 1.99±0.23x $10^6$ |
| | Viability (%±SD) | 90.8±6.7 | 86.2±8.1 | 91.1±7.4 | 88.4±7.5 |
| 6-8 | Cell yield (No.±SD) | 31.8±7.7x $10^6$ | 892.1±69x $10^3$ | 151.0±53x $10^4$ | 2.22±0.30x $10^6$ |
| | Viability (%±SD) | 95.2±7.1 | 89.7±8.1 | 92.6±7.4 | 89.9±6.6 |
| ≥18 | Cell yield (No.±SD) | 10.7±6.2x $10^6$ | 309±175x $10^3$ | 97.3±16.5 x$10^4$ | 1.20±0.13x $10^6$ |
| | Viability (%±SD) | 86.2±7.4 | 85.6±6.2 | 88.2±8.9 | 86.8±9.8 |

**C**

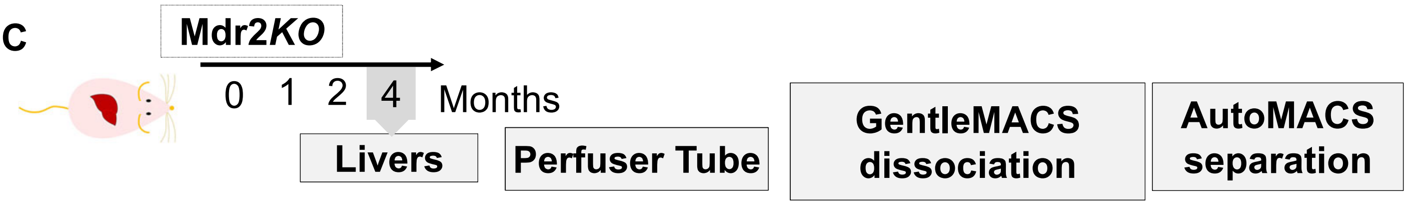


**D**

| | HC | HSC | KC | LSEC |
|---|---|---|---|---|
| Cell yield (No.±SD) | 4.5x$10^{5\pm}$2.09 | 100±50x$10^3$ | 2.8±0.82x$10^6$ | 2.0±0.27x$10^6$ |
| Viability (%±SD) | 85,9 ± 8.2 | 88.8±9.2 | 89.2±9.8 | 87.9±11.3 |

**E**

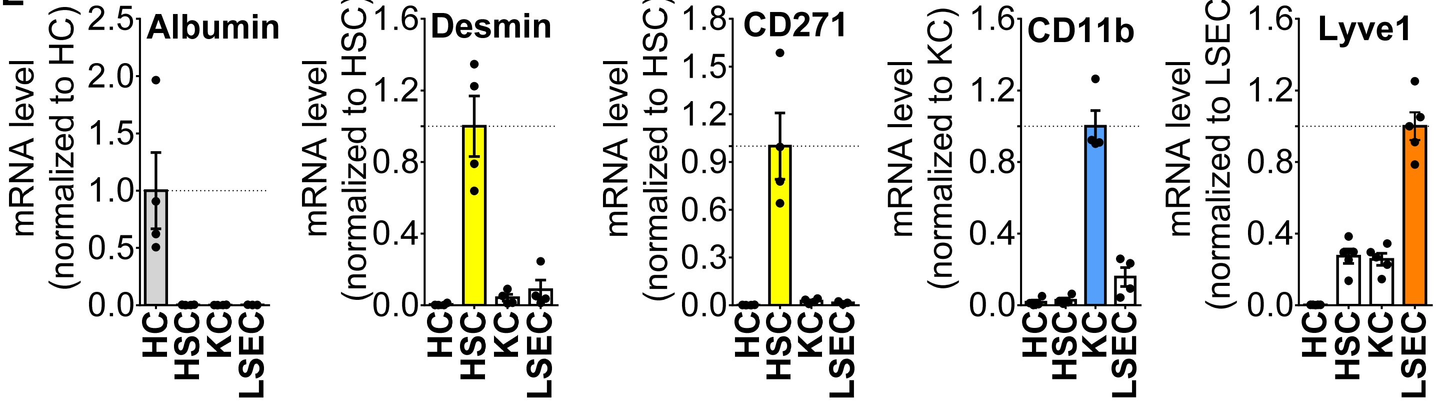
