## Supplementary figures and images for "CD271 sorting for improved liver cell isolation: Semiautomated and simultaneous preparation of parenchymal and non-parenchymal cells from mouse and human livers"

### Graphical Abstract

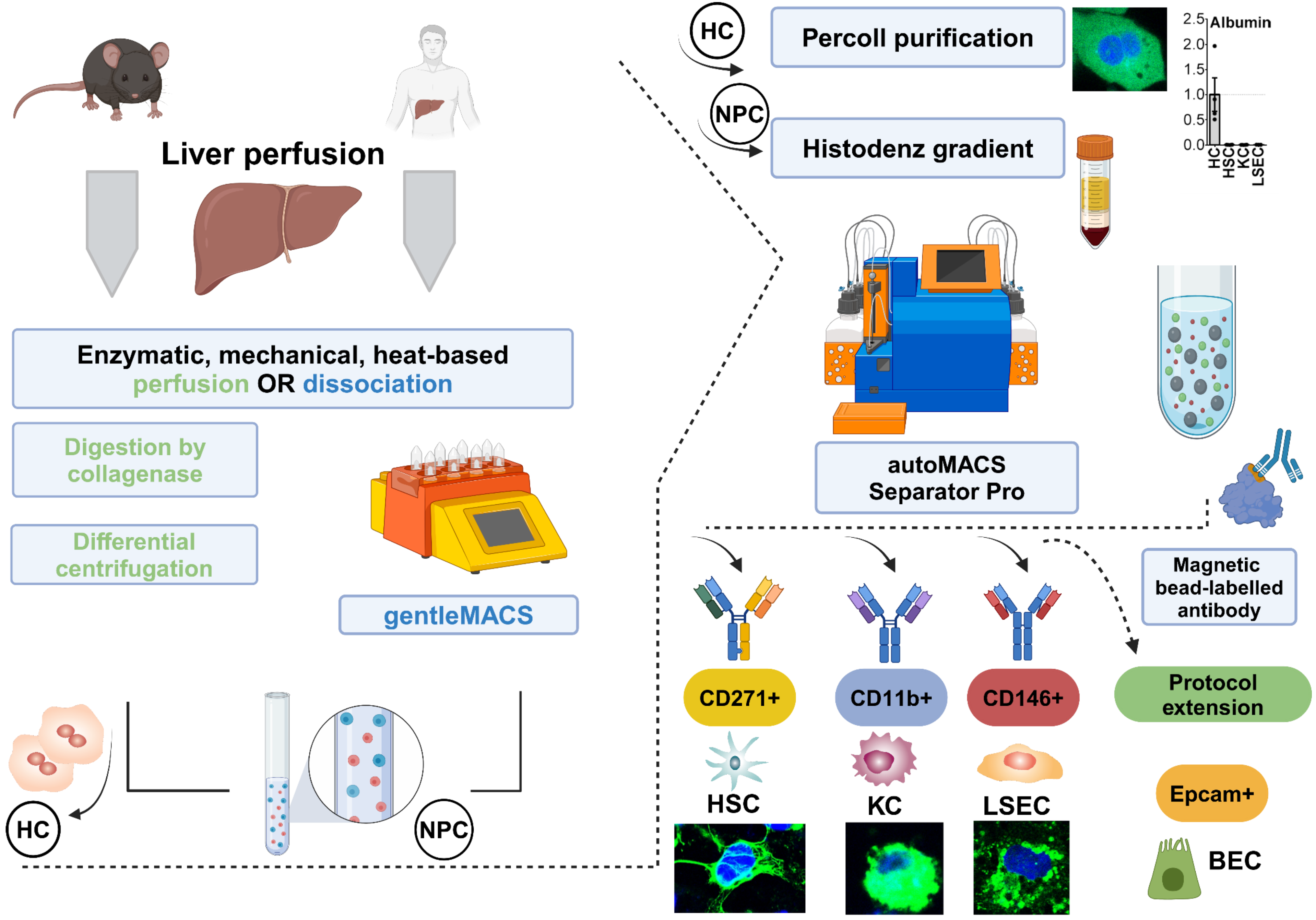
